# Novel transient cell clusters provide a possible link between early neural activity and angiogenesis in the neonatal mouse retina

**DOI:** 10.1101/2022.08.04.502860

**Authors:** Jean de Montigny, Courtney Thorne, Diya Bhattacharya, Dimitrios Bousoulas Sertedakis, Vidhyasankar Krishnamoorthy, Fernando Rozenblit, Tim Gollisch, Evelyne Sernagor

## Abstract

Developing neurons become spontaneously active while growing blood vessels begin to irrigate their surroundings. However, surprising little is known about early interactions between neural activity and angiogenesis. In the neonatal mouse retina, spontaneous waves of impulses sweep across the ganglion cell layer (GCL), just underneath the growing superficial vascular plexus. We discovered clusters of transient auto-fluorescent cells in the GCL, forming an annulus that co-localizes with the frontline of the growing plexus. Blood vessel density is highest within cluster areas, suggesting their involvement in angiogenesis. Once the clusters and blood vessels reach the retinal periphery by the end of the first postnatal week, the clusters disappear, eliminated by microglial phagocytosis. Electrical imaging suggests that they have their own electrophysiological signature. Blocking Pannexin1 (PANX1) hemi-channels with probenecid blocks the waves and the fluorescent clusters disappear following prolonged exposure to the drug. Spontaneous waves’ initiation points follow a developmental center-to-periphery progression similar to the cluster cells. We suggest that these transient cells are specialized, hyperactive neurons residing in the GCL. They generate spontaneous activity hotspots, thereby triggering waves through purinergic paracrine signaling via PANX1 hemi-channels. The strong activity generated around these hotspots triggers angiogenesis, attracting new blood vessels that provide local oxygen supply. Signaling through PANX-1 attracts microglia that establish contact with these cells, eventually leading to their elimination by phagocytosis. These cluster cells may provide the first evidence that specialized transient neuronal populations guide angiogenesis in the developing CNS through neural activity.

## Introduction

By continuously producing electrical signals, neurons are amongst the most energy-demanding cells in the organism. Resting ionic levels are restored by metabolic pumps through oxygen supplied by blood vessels via the neurovascular unit. Intense spontaneous neural activity is omnipresent in the developing CNS (Blankenship and Feller, 2010). It occurs during short, well-defined periods, coinciding precisely with blood vessels initial irrigation of neural territories (Paredes et al., 2018). Such coincidence may reflect a universal mechanism underlying CNS angiogenesis. A plethora of angiogenic signaling molecules have been identified, but surprisingly little is known about the role of neural activity *per se* in guiding angiogenesis (Andreone et al., 2015; Paredes et a;. 2018). This is partly due to the challenge of investigating developing brain neurovascular networks in tri-dimensional space.

With its planar organization, the retina offers unique opportunities to study these important questions. In mouse, the primary superficial plexus emerges from the optic nerve head (ONH) shortly after birth and expands just above the ganglion cell layer (GCL) in two-dimensional fashion, reaching the periphery by postnatal day (P) 7 (Paredes et al., 2018).

While the superficial plexus expands from the ONH to the periphery, waves of intense bursting activity spread across the GCL (Meister et al, 1991). Profound changes in wave spatiotemporal features occur with development (Maccione et al, 2014), reflecting changes in the cellular mechanism underlying their generation/propagation. In mouse, it switches from gap junction communication (Stage-1, prenatal) to cholinergic neurotransmission originating from starburst amacrine cells (SACs) (Stage-2, P0-10) (Feller et al, 1996), and finally, to glutamatergic bipolar cells before waves disappear around eye opening (Stage-3, P10-13).

Completion of the superficial plexus outgrowth coincides with the disappearance of Stage2 waves, suggesting a possible causal link between these events. However, specifically manipulating cholinergic activity mostly affects the deep plexus formation during the second postnatal week (Weiner et al, 2019), indicating that cholinergic signaling *per se* does not mediate the earliest angiogenesis phase, despite being involved in blood flow control in the adult retina (Ivanova et al., 2016).

Here we report for the first time that transient clusters of cells are present on the edge of the superficial vascular plexus in the GCL during the period of Stage-2 waves. We propose that these cells are specialized neurons that guide angiogenesis by generating activity hotspots that trigger wave initiation through paracrine signaling.

## Results and Discussion

While exploring the expression of various markers during Stage-2 waves, we serendipitously discovered sparse auto-fluorescent cellular clusters (Fig1A) in the GCL (Fig1E,F). Cluster cells are larger than retinal ganglion cells (RGCs) and SACs. Retinal sections reveal that they are in close proximity to SACs and may even contact them (Fig1F, arrow). They auto-fluoresce over a broad wavelength spectrum (Fig1D) in fixed and live retinas. They form a small annulus around the ONH from P2, gradually expanding and, reaching the periphery by P7 (Fig1A-C). The most substantial changes occur between P3-5.

**Figure 1:**
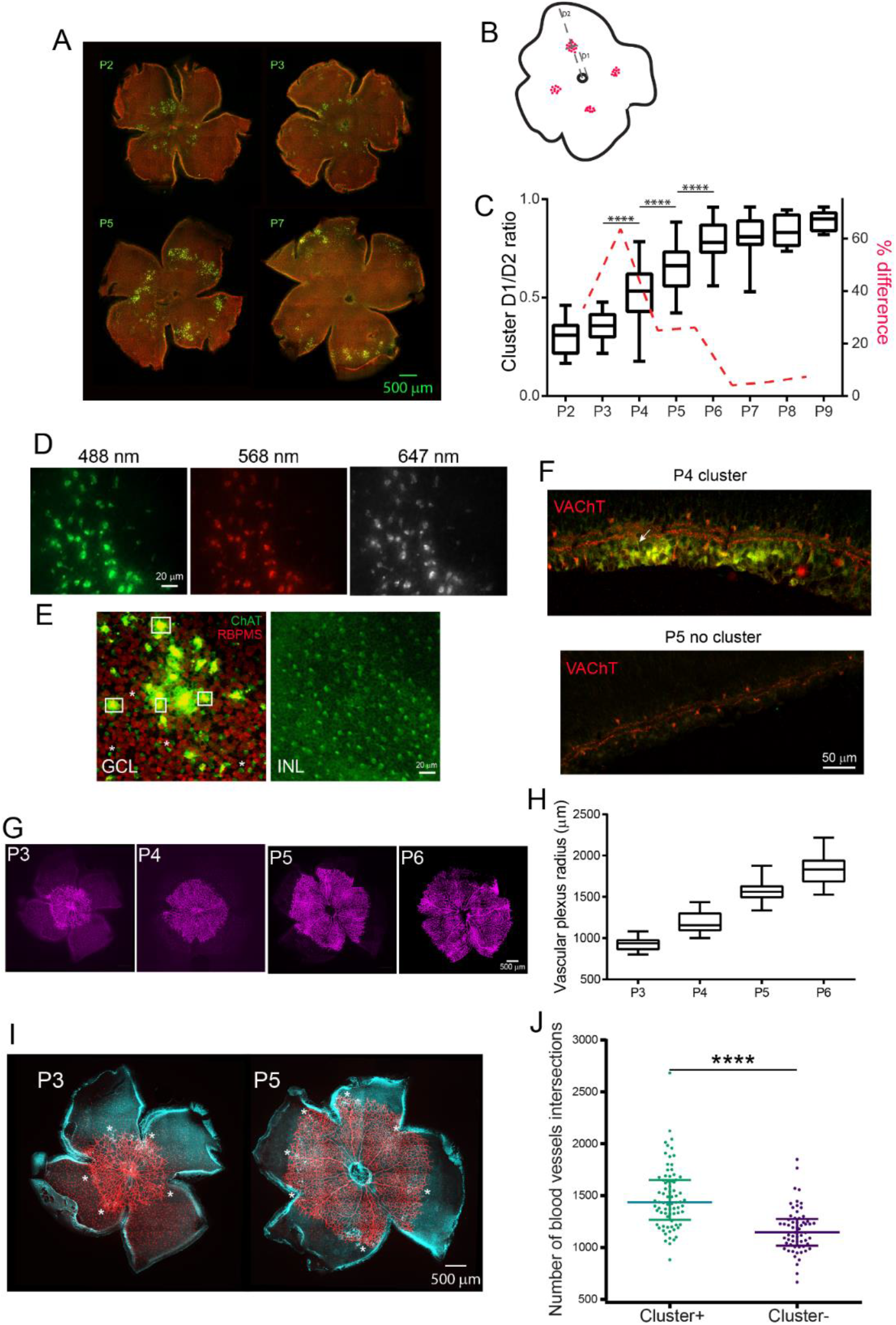
Transient auto-fluorescent cell clusters in the neonatal mouse retina. **A**: Mouse retinal wholemounts stained for ChAT (green) and RBPMS (red). Scale bar: 500 µm. **B**: Method for calculating the relative position of clusters between the ONH (small black circle in the middle) and periphery. Cluster cells are represented by red dots. D1: distance from center of ONH to center of cluster. D2: distance between center of cluster to periphery. **C**: Box plot showing developmental changes in D1/D2 ratios. Each box illustrates the median (horizontal line) and interquartile range, with minimum and maximum values (whiskers). Asterisks indicate significant changes between consecutive days (One-way ANOVA with post-hoc Tukey test). The red dotted line illustrates the percentage difference in values between consecutive days, showing peak difference between P3 and P4 and no further changes from P6 onwards. **D**: One auto-fluorescent cell cluster (P5 retina, fixed retina) visualized at three different wavelengths without any immunolabeling, indicating intrinsic fluorescent signals. **E**: P5 cluster viewed at the GCL level and at the inner nuclear layer (INL) level. Green: ChAT; red: RBPMS. The cluster cells are visible only at the GCL level (examples in white boxes) and they are much larger than SACs (white asterisks) or RGCs (in red). At the INL level, there are only SACs. **F**: Retinal sections showing VAChT (red) expression in a cluster area (P4 retina) and in an area devoid of clusters (P5 retina). SACs express VAChT, exhibiting the typical double laminar expression in the IPL flanked by cell bodies in the INL and GCL. In areas devoid of clusters, only the SAC expression pattern can be seen. **G**: The superior vascular plexus (labeled with isolectin B4 (IB4)) grows from the ONH, where it emerges shortly after birth, and reaches the periphery by the end of the first postnatal week. Shown at P3-4-5-6. **H**: Box plot (median with interquartile range, minimum and maximum values (whiskers)) showing ongoing expansion of the superior vascular plexus between P2-6. **I**: the auto-fluorescent cluster cells (white) are located under the front edge of the vascular plexus (IB4, red). Clusters are indicated with white asterisks (P3 and P5 retina). Microglia are well expressed with IB4 too in these retinas. **J**: Scatter plot of blood vessel branch intersections (Sholl analysis). There are significantly more branches in cluster areas (cluster +, green) than in clusterless areas (cluster -, purple). Median with interquartile range. ****: P<0.0001, Mann Whitney 2-tailed test.

The clusters begin to disintegrate by P7-8 and completely disappear by P10, coinciding with the switch from Stage-2 to Stage-3 waves.

The superficial vascular plexus gradually expands above the RGC layer (Fig1G,H), reaching the periphery by P7-8 as well. The cluster cells are always positioned underneath the front edge of the growing vasculature, but never ahead of it, regardless of the developmental stage (Fig1I, asterisks), suggesting that they may play an angiogenic role in the growing plexus. Sholl analysis (see Methods) indeed reveals significantly more blood vessel branches in areas with clusters than in areas without clusters at matching eccentricities (Fig1J).

Their location in the GCL and co-localization with the front edge of the primary plexus suggest that the cluster cells may be a specialized, transient RGC type involved in angiogenesis. In support, the primary plexus does not develop in the absence of RGCs (Edwards et al., 2012; Kim et al. 2011).

Sparsity may be the reason these cells were never reported in previous studies. Only a few clusters are visible in each retina, necessitating pan-retinal visualization for reliable detection. Moreover, their auto-fluorescence warrants that they are always visible, even with markers for other cell types, preventing reliable identification. Other approaches such as single-cell RNA sequencing and non-fluorescent immunolabeling are thus necessary to establish their genetic identity.

The cell clusters are present only during Stage-2 waves, which made us wonder whether they may be involved in their generation. To explore this possibility, we recorded waves between P2-13 using large-scale multielectrode arrays (MEAs) with 4,096 active electrodes spanning an area of 5.12×5.12 mm^2^, electrode pitch 81 µm, large enough to cover the entire retinal surface at all ages (Fig2). Wave origins (determined as the xy coordinates of the initial wave center of activity trajectory, see Maccione et al., 2014) were aligned with the image of the retina itself (Fig2A) and classified as either central (C) or peripheral (P) (Fig2B). Fig2C reveals a very clear centrifugal developmental shift in wave origins (with P/C ratio values reaching >1), with peak changes between P3-5, exactly like for the cluster locations (Fig1C). That trend disappears and P/C ratios drop to their lowest values once waves become driven by glutamate (Stage-3), characterized by small activity hotspots that tile the retina (Maccione et al., 2014). When xy wave origin values are randomly shuffled between P and C, the developmental centrifugal trend disappears, with P/C ratios approaching 1 at all ages (Fig2C, red). In stark contrast with all previous studies on retinal wave dynamics, our findings thus suggest that during Stage-2, wave initiation is highly non-random, following the same centrifugal trajectory as the cluster cells and vascular plexus outgrowth. We previously reported that waves grow until P6 (Maccione et al, 2014). This may be due to their initiation points becoming gradually more peripheral. Altogether, our observations raise the possibility that the cluster cells act as “pacemakers”, triggering wave initiation in gradually more peripheral hotspots. Once triggered, waves spread across the RGC layer via the horizontal SAC cholinergic network (Butts et al., 1999; Hennig et al., 2009).

**Figure 2:**
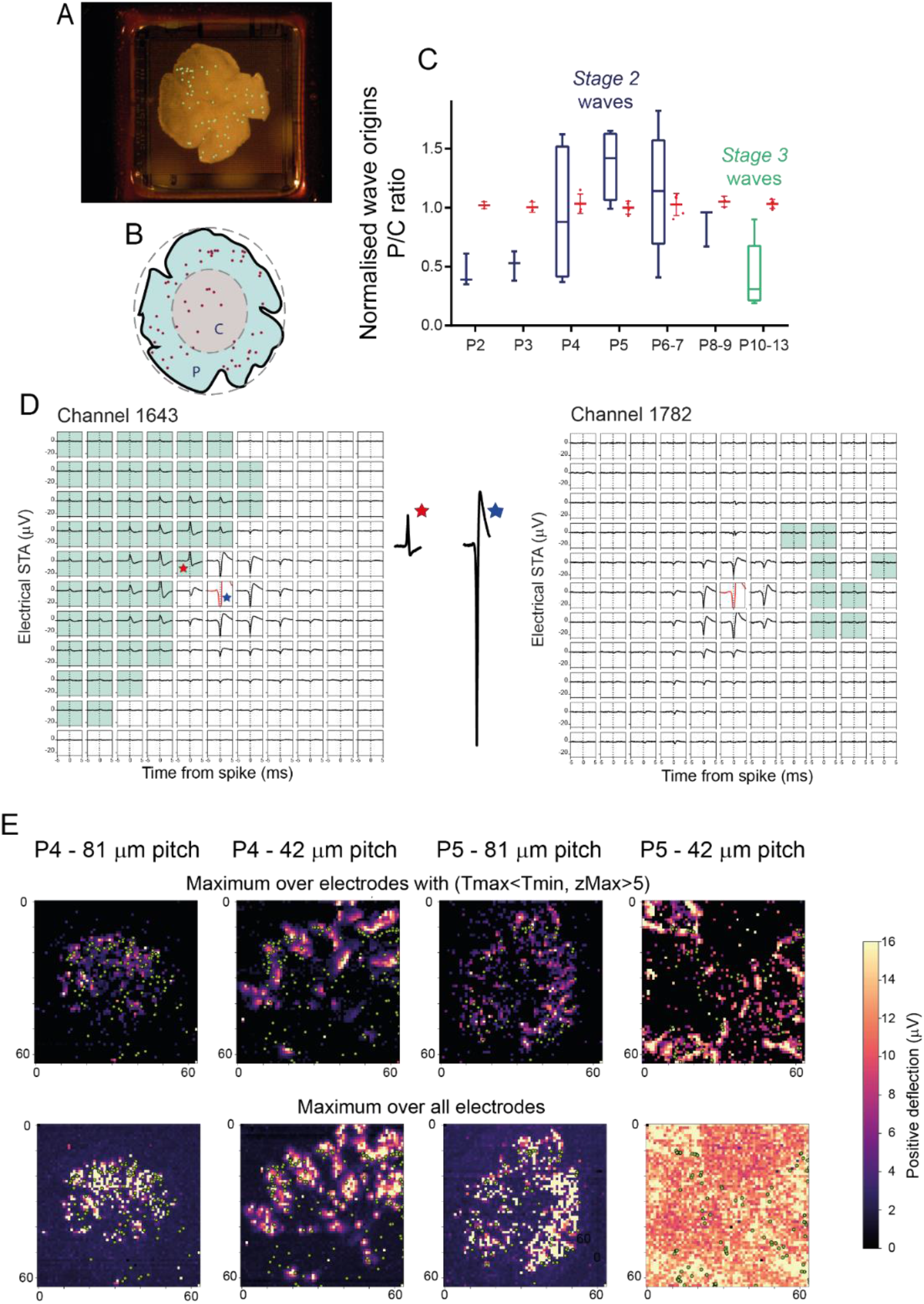
retinal waves and cluster cells. **A**: Photograph of a P4 retina on the MEA, taken at the end of the recording session. Wave origins (green dots) are overlaid on the photograph. **B**: Method to measure wave origins in center and periphery. P: periphery, C: center. Red dots: wave origins. **C**: box plot illustrating the P/C ratios from P2-13. Wave origins expand from center to periphery between P2 and P5-6, similar to the clusters themselves (Fig1C). Same boxplot conventions as for Fig1C. Statistical analysis was not possible here due to the small numbers of values in each group (one ratio value per retina). Blue: Stage-2 cholinergic waves; green: Stage-3 glutamatergic waves; red: Monte Carlo randomized P/C ratio values. **D**: Electrical imaging of retinal waves. STAs for two trigger channels showing signals averaged over -5 to +5 ms relative to spikes on the trigger channel (in red) for 11×11 surrounding recording channels. Recording channels with dipole activity (with maximal positive deflections with z score>5 occurring before the maximal negative deflection, see Methods) are marked with the green mask. Channel 1643 has a marked area with dipole signals near the trigger channel. The single traces on the right side of the electrode grid show the full size of the spike on the trigger channel (blue asterisk) and maximal positive deflection (red asterisk), emphasizing the fact that the amplitude of the dipole signals is significantly smaller than spikes, suggesting that they may represent slow, paracrine signals. (P4 retina, 60 min recording). **E**: Maximal projections for signals with pre-STA spike signals with larger positive than negative deflections (top row) and for all maximal projections (bottom row) over the entire MEA. Maps are shown for two P4 and two P5 retinas, each with one example recorded on an array with 42 µm electrode pitch, and for another array with 81 µm pitch. Wave origins (green dots) are overlaid on the electrical signals.

We used electrical imaging (Greschner et al, 2016; Zeck et al., 2011; Petrusca et al., 2007; Litke et al, 2004) (see Methods) to visualize wave-related electrical activity at high spatiotemporal resolution (Fig2D,E). We used MEAs with either 81 or 42 µm electrode pitch. The latter, albeit covering a smaller area, provide higher spatial resolution. Negative deflections smaller than the spike used for spike-triggered averaging (STA) (Fig2D, blue asterisk) were detected around most STA channels, presumably reflecting wave-related activity propagation (Channel 1782, Fig2D). But for some channels there were smaller (yet significantly distinguishable from baseline) positive signals (Fig2D, red asterisk) emerging simultaneously with the negative spike signal (Channel 1643, Fig2D). Overall, when combining the activity footprint from all channels exhibiting such positive-negative “dipole” behavior (Fig2E), we found that these areas form clusters in close proximity with wave origins (Fig2E, green dots), suggesting that these signals may reflect activity originating from the transient cell clusters. However, when plotting all maximal projections, regardless of whether they have a positive deflection or not (Fig2E, bottom row), co-localization with the wave origins is less conspicuous.

These small dipole signals look different from “classical” somatic spikes (like the STA spikes), and may reflect a distinct type of electrical activity, perhaps akin to slow, paracrine signaling (see below).

The auto-fluorescent cluster cells are always underneath the expanding plexus frontline, progressively disappearing from more central areas. We found that they disappear through microglial phagocytosis under the expanding plexus frontline (Fig3A), with microglia exhibiting distinct activated morphology (shorter and thicker processes) around them. As a matter of fact, the cluster cells appear to attract microglia, indicated by a positive correlation between cluster cells and microglia densities (Fig3C). Microglial density is significantly higher within cluster areas than in areas without clusters at matching eccentricities (Fig.3B), whilst there is no difference in areas ahead or behind the clusters (Fig3D,E).

**Figure 3:**
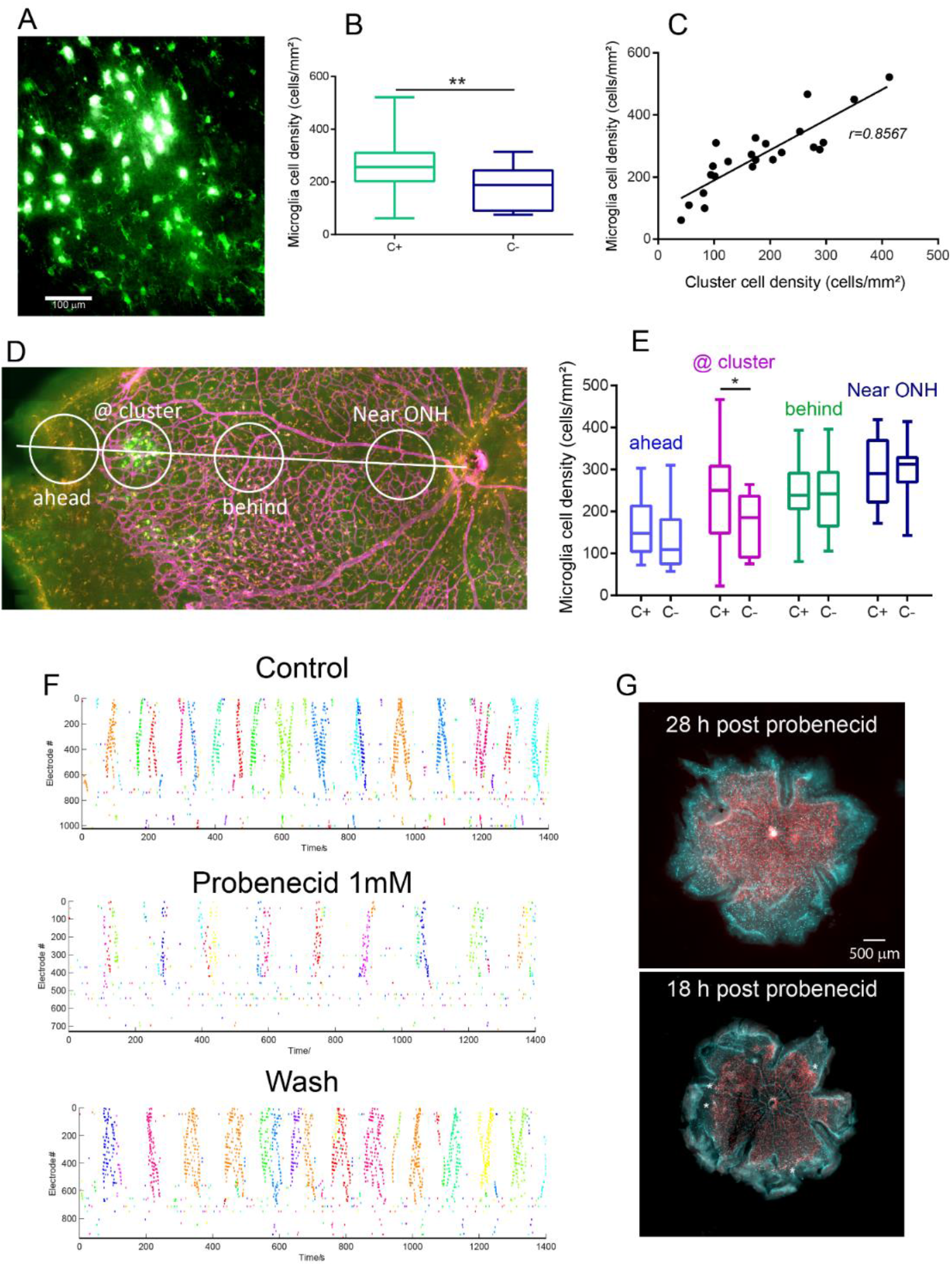
Cluster cells and microglia. **A**: Micrograph of auto-fluorescent cluster cells (white) surrounded by microglial cells labeled with iba1 (green). Resting microglia have thinner and more numerous processes than once they are activated and encircle the cluster cells. **B**: Box plot (median and interquartile ranges) comparing microglia density within cluster areas (C+) versus clusterless areas (C-). P<0.0138. Mann Whitney two-tailed test. **C**: Strong positive correlation between the densities of microglial cells and auto-fluorescent cluster cells (p<0.0001). **D**: Method for measuring microglia density at different retinal eccentricities, with measures taken near the ONH, between the ONH and clusters (or clusterless areas at corresponding eccentricities), at clusters (or corresponding clusterless areas) and ahead of clusters, further in the periphery, in the unvascularized part of the retina. The micrograph shows blood vessels labeled with IB4 (magenta), microglia labeled with iba1 (orange) and auto-fluorescent cluster cells (white). **E**: box plot (with medians and interquartile ranges) showing that only the density of microglia is significantly higher only within clusters (p<0.01, Mann Whitney, two-tailed test). **F**: Raster plots of waves in control conditions, in the presence of probenecid 1mM (recording started 30 minutes after adding the drug), and immediately following washout. Each row represents an active electrode on the MEA. Waves are indicated by synchronized activity across channels. The activity is shown against time (in seconds). Waves are detected according to methods described in Maccione et al., 2014. Each detected wave is in a different color. Probenecid profoundly reduces retinal excitability, with significant decrease in wave frequency and number of channels recruited within waves. **G**: Retinal wholemounts stained for blood vessels (IB4, red) and microglia (iba1, cyan) following overnight exposure to probenecid. Cluster cells are barely visible in the bottom retina (white asterisks), 18 hours following exposure to probenecid, and are not visible in the top retina, following 28 hours in probenecid.

Neural activity triggers microglia activation and contact with neurons (Fontainas et al., 2011; Li et al., 2012). The process is mediated via voltage-gated pannexin-1 (PANX1) hemi-channels that release ATP acting on P2 purinergic receptors located on microglia (Velasquez and Eugenin, 2014), sending an “eat me” signal to microglia. In the retina, PANX1 hemi-channels are particularly prominent on RGCs (Dvoriantchikova et al., 2018). To verify whether the cluster cells may use that mechanism, we have investigated the effect of probenecid, a PANX1 inhibitor (Silverman et al., 2008; Dvoriantchikova et al., 2018).

Probenecid (1mM) had a powerful and immediate inhibitory effect on wave generation, leading to strong decrease in wave frequency and cellular recruitment within waves (Fig3F). Following overnight incubation in probenecid, clusters were either not visible or much smaller (Fig3G) and waves disappeared. These observations indicate that signaling through PANX1 hemi-channels controls retinal excitability, corroborating one earlier study showing that cyclic-AMP signaling controls retinal wave dynamics (Stellwagen et al., 1999).

Disappearance of the fluorescent cluster cells following overnight exposure to probenecid suggests that they become fluorescent only upon contact with microglia, perhaps because of metabolic stress (e.g. lipofuscin expression; Wilhelm et al., 2011). Perhaps these same cells are initially non-fluorescent, located just ahead of the vasculature front while they are most active in terms of triggering waves and attracting blood vessels. Calcium imaging approaches will help us decipher these questions.

In summary, these cluster cells may provide the first evidence that specialized transient neuronal populations guide angiogenesis in the developing CNS through neural activity.

## Methods

### Animals

All experimental procedures were approved by the UK Home Office, Animals (Scientific procedures) Act 1986.

Experiments were performed on neonatal (P2-13) C57bl/6 mice. All animals were killed by cervical dislocation followed by enucleation.

### Immunostaining and imaging

For our anatomical studies, retinas were extracted from pups aged P2 (N=18 retinas), P3 (N=19), P4 (N=23), P5 (N=25), P6 (N=16), P7 (N=13), P8 (N=10), P9 (N=10), P10 (N=4) and P11 (N=4).

We have used the following antibodies:

### Primary antibodies

ChAT (AB144P, goat polyclonal, Merck Millipore).

VAChT (PA5-77386, rabbit polyclonal, ThermoFisher Scientific).

RBPMS (1830-RBPMS, rabbit polyclonal, Phosphosolutions), against RGCs (Rodriguez et al., 2014).

Iba1 (019-19741, rabbit polyclonal, Alpha Labs)

### Secondary antibodies

Donkey anti rabbit Alexa 568 (A10042, Invitrogen).

Donkey anti goat Dylight 488 (SA5-10086, Thermo Fisher Scientific). Goat anti-rabbit A546 (A11010, Invitrogen)

### Blood vessels staining

Isolectin B4 fluorescein (FL-1201, Vector Labs) Isolectin B4 Dylight 594 (DL-1207, Vecor Labs) Isolectin B4 DyLight649 (DL-1208, Vector Labs)

### Retinal sections

Eyecups were prepared from mouse pups aged P2-P9, fixed for 45 min in 4% paraformaldehyde (PFA), incubated in 30% sucrose in 0.1M phosphate buffer solution (PBS) for at least 12 hours, and then embedded in Optimal Cutting Temperature (OCT) cryo embedding compound and frozen at -20°C. Eyecups were sliced as 28μm thick sections using a cryostat (Model: OTF5000, Bright Instruments), washed with PBS to remove OCT, and incubated in blocking solution for 1 hour (5% secondary antibody host species serum with 0.5% Triton X-100 in PBS) prior to staining with antibodies.

Retinal sections were incubated with the primary antibody solution (0.5% Triton X-100 with VAChT (1:500 in PBS)) for 12 hours at 4°C. Sections were washed with PBS, followed by incubation with fluorescent secondary antibody solution (0.5% Triton X-100 with donkey anti rabbit Alexa 568 (1:500 in PBS) for 1 hour.

Finally, slices were washed with PBS and embedded with home-made OPTIClear refractive-index homogenisation solution. OPTIClear solution consists of 20% w/v N-methylglucamine, 25% w/v 2,2’-Thiodiethanol, 32% w/v Iohexol, pH 7-8. The solution is clear and colourless, with a refractive index of 1.47-1.48.

Sections were imaged using the Zeiss LSM 800 confocal microscope. Regions of interest were selected by focusing on clusters.

### Retinal wholemounts

Wholemount retinas were prepared from mouse pups aged P2-P11, flattened on nitrocellulose membrane filters and fixed for 45 min in 4% PFA. Retinas were then incubated in blocking solution (5% secondary antibody host species serum with 0.5% Triton X-100 in PBS) for 1 hour.

Primary antibodies: 0.5% Triton X-100 with RBPMS (1:500), ChAT (1:500), iba1 (1:1000).

Secondary antibodies: 0.5% Triton X-100 with donkey anti rabbit Alexa 568 (1:500), donkey anti goat Dylight 488 (1:500), goat anti-rabbit A546 (1:500).

Isolectin B4 (IB4) staining of blood vessels (1:250) was performed together with the secondary antibody staining step.

Retinas were incubated with the primary antibody solution for 3 days at 4°C, then washed with PBS and incubated with the secondary antibody solution for 1 day at 4°C (with or without IB4).

Finally, retinas were washed with PBS and embedded with OptiClear.

Zeiss AxioImager with Apotome processing and the Zeiss LSM 800 confocal microscope were used to image the retinas.

High-resolution images of the RGC layer down to the INL were obtained by subdividing retinal wholemounts into adjacent smaller images that were subsequently stitched back together to view the entire retinal surface. Regions of interests were selected around the clusters.

To compensate for variability in retinal thickness, several focus points were set across the retinal surface in order to keep sharp focus on the desired cell layer. Each individual picture was then acquired in all color channels at 20x magnification, and with 10% overlap between neighboring areas. This overlap is used to correctly align and stitch together all pictures using the Zen Pro software (Zeiss). Z-stacks of images at 40x magnification were acquired at regions of interest to visualize cells in 3D. Z-stacks consisted of images taken every 1 μm from the RGC layer to below the INL.

Some wide field fluorescent images were acquired with the Nikon Ni-E microscope. Multiple images were taken at 10x or 20x, and then combined in order to obtain a high-resolution image of the whole retina.

### Data processing and analysis

To calculate the relative position of the cell clusters (see Fig1B) between the ONH and periphery, lines were traced and measured from the middle of the ONH to the middle of a cluster (D1) and then from the same point in the cluster to the periphery of the retina (D2). D1/D2 represents the relative position of the clusters. One-way ANOVA was used on all 233 ratio values for all eight groups. Tukey post-hoc test was used to identify significant changes in cluster positions between consecutive developmental days.

Sholl analysis was performed using Fiji to quantify blood vessel branches. IB4 stained vasculature images from retinal wholemounts were rendered binary using the ‘Threshold’ function. To remove signals inherent to auto-fluorescence of the cluster cells, a separate color channel image showing only cluster cells was rendered binary and subtracted from the vasculature image. Subtracted images were then adjusted using the ‘Brightness/Contrast’ function to obtain solid binary images.

To quantify blood vessel density in the vicinity of clusters, segmental sections of the binary, cluster-subtracted whole retina vasculature images were outlined using the ‘Angle’ tool (allowing measurement of the angle of the segment and subsequent calculation of the surface area of the region of interest (ROI)) either with or without a cluster at the edge of the vascular plexus. Sholl rings were set from the center of the ONH with a step size of 30 µm. Measurements were not taken from the inner 50 % of the segment (based on the radius of the vascular plexus at its largest point). Intersections were counted manually.

To quantify blood vessel branch density in areas with clusters versus areas without clusters at matching eccentricities, ROIs were outlined using the ‘Polygons’ function either around clusters (Cluster+ ROI), or in adjacent clusterless regions at similar eccentricities (Cluster-ROI). This was done in images from a color channel in which the blood vessels were not visible. Once all ROIs were outlined for one retina, they were overlaid on the binary, cluster subtracted vasculature image for that same retina. Any ROIs noticeably outside of the vascular plexus were discounted. Sholl rings were set from the center of the ONH with a step size of 30 µm. Intersections were counted manually.

All cell counts were performed using FIJI. Cluster cells were counted manually in retinal wholemount images using the ‘Multi-point’ function. Cells were identified by auto-fluorescence using an image captured on a channel without any fluorescing immunostaining. The vascular plexus was outlined using the ‘Freehand’ tool and ‘Measure’ function was used to determine surface area. Clusters were delimited and cells counted manually, identified by auto-fluorescence. A ‘cluster’ was defined as an uninterrupted group of auto-fluorescent cells spatially separated from any other group, localised near the outer edge of the superficial vascular plexus. No lower or upper size limit was assigned to clusters.

To quantify microglia and cluster cell densities, radial straight lines were drawn from the center of the ONH to the periphery of the retina, either through the centre of a cluster or through an adjacent clusterless region. This was only performed where radial lines reached the periphery of the retina, uninterrupted by cuts. ROIs were outlined using the ‘Oval’ selection tool at four points along each radial line to comprise an ROI set. The four ROI types in each set were; ahead of the cluster/clusterless region, at the cluster/clusterless region, behind the cluster/clusterless region, and close to the ONH (Fig3D). ROIs were outlined at comparable eccentricities for each set. Iba1 stained microglia were counted manually. All Iba1 stained whole retinas were from P5 or P6 animals. Cluster cells were identified by auto-fluorescence and counted manually.

### Electrophysiology-MEA recordings

Retinas were isolated from mouse pups at P2 (N=4 retinas), P3 (N=4), P4 (N=10), P5 (N=14), P6 (N=3), P7 (N=2), P8 (N=2), P9 (N=2), P10 (N=2), P11 (N=2), P12 (N=1), P13 (N=1). The isolated retina was placed, RGC layer facing down, onto the MEA and maintained stable by placing a small piece of polyester membrane filter (Sterlitech, Kent, WA, USA) on the retina followed by a home-made anchor. The retina was kept in constant darkness at 32°C with an in-line heater (Warner Instruments, Hamden, CT, USA) and continuously perfused using a peristaltic pump (∼1 ml/min) with artificial cerebrospinal fluid containing the following (in mM): 118 NaCl, 25 NaHCO_3_, 1 NaH_2_PO_4_, 3 KCl, 1 MgCl_2_, 2 CaCl_2_, and 10 glucose, equilibrated with 95% O_2_ and 5% CO_2_. Retinas were allowed to settle for 2 hours before recording, allowing sufficient time for spontaneous activity to reach steady-state levels. Probenecid (4-(dipropylsulfamoyl)benzoic acid) (water soluble form, ThermoFisher Scientific, P36400) was used at 1mM (after calibrating the effect between 250 µM and 1 mM).

High resolution extracellular recordings of spontaneous waves were performed as described in details in Maccione et al. (2014), using the BioCam4096 platform with APS MEA chips type HD-MEA Stimulo (3Brain GmbH, Switzerland), providing 4096 square microelectrodes of 21 μm x 21 μm in size on an active area of 5.12 × 5.12 mm^2^, with an electrode pitch of 81 μm. Two P5 and one P4 datasets were acquired with the MEA chip HD-MEA Arena (2.67×2.67mm^2^ active area, electrode pitch 42 μm) for electrical imaging.

Raw signals were visualized and recorded at 7 kHz sampling rate with BrainWaveX (3Brain GmbH, Switzerland). Each dataset consisted of 30 min of continuous recording of retinal waves. The datasets used for electrical imaging were acquired at 17.855 kHz for 30 or 60 min.

In the BioCam4096, samples of MEA signal are acquired row-wise by the amplifier. Individual samples consist of 64 columns and 64 rows and often show a small but measurable bias across rows of ca. 2-4 μV (1-2 ADC units). While such bias is negligible for most applications, it does degrade the quality of electrical images. Therefore, to reduce the bias before electrical imaging, the median value of each row was independently calculated and subtracted.

Retinas were photographed on the MEA at the end of the recording session to ensure we document the precise orientation of the retina with respect to the array of electrodes (Fig2A,B).

### Data processing and analysis

Burst and wave detection was done in Matlab (Mathworks) as described in Maccione et al. (2014). We computed the coordinates of the wave origins (Maccione et al., 2014) and aligned them with a picture of the retina on the MEA (Fig2A). Retinas were delimited by a peripheral annulus (P), surrounding a concentric central ellipse, 50% smaller (C) (Fig2C). We counted wave origins (Fig2C, red dots) localized in C (grey shading) and in the retinal area confined within P (green shading) and plotted the P/C ratio between P2-13 (Fig2D). For Monte Carlo randomization of wave origins, the process was repeated ten times.

Electrical images were computed independently for each electrode by averaging the electrical activity in the MEA surrounding the time of spikes in that electrode (spike-triggered average). First, spikes were detected independently at each electrode with the default detection parameters of BrainWaveX (3Brain GmbH, Switzerland) (Maccione et al., 2014). To ignore electrodes without good physical coupling to the retina, only active electrodes – those with any noteworthy activity – were analyzed. Active electrodes were defined as having a normalized spike count

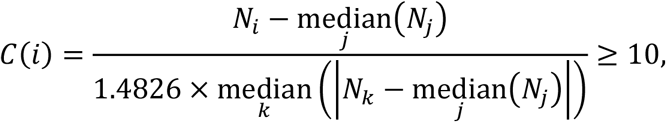

where *N* stands for the spike count from an electrode. The denominator of *C* robustly estimates the standard deviation of spike counts using the median absolute deviation from the median (Quiroga et al., 2004; Donoho & Johnstone, 1994).

The electrical imaging itself proceeded as follows. For each active electrode, snippets of raw MEA signals were taken from -5 to +5 ms of the detected spike times. Only spikes appearing concomitantly with retinal waves were considered (wave onset and offset were determined by (i) detecting bursts on each electrode separately, and (ii) grouping bursts into waves based on temporal overlap and proximity, see Maccione et al., 2014 for further details). For the sampling rate of 17.855 kHz, a snippet consisted of 64 rows, 64 columns, and 180 sample points. Averaging the snippets thus led to a 64×64×180 movie *A*_*i*_(*x, y, t*) representing the typical electrical activity in the temporal vicinity of spikes from electrode *i*. To remove noisy electrodes, electrical images whose negative peak of the spike had a half-peak width of less than 0.3 ms were discarded from the remaining analysis. To visualize the electrical activity, movies *A*_*i*_(*x, y, t*) were reduced to a map of negative deflections 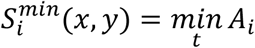, positive deflections 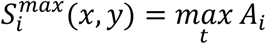, and the times 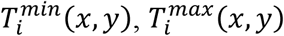 where such deflections occurred.

In some electrical images, a dipole activity was observed where positive deflections emerged simultaneously with the expected negative deflections of a spike. These positive lobes showed a strong positive deflection followed by a negative deflection (Fig2D). To detect candidate regions with positive lobes, pixels (x,y) were selected when they had a significant positive deflection at a time 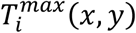 preceding the time of negative deflection 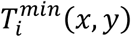 and not occurring later than 0.5 ms after the triggering spike. The latter condition helped to remove pixels exhibiting purely axonal propagation. Positive deflections were considered significant when their z-scored values

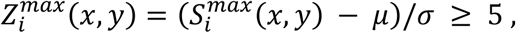

with 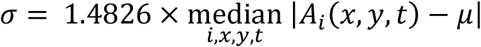 and 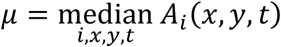 (see C(i) above).

Defining a mask *W*_*i*_(*x, y*) as unity for such selected pixels with positive deflections (green background in Fig2D) and zero everywhere else, the maximum projection (Fig2E) over selected regions was given by 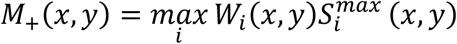. The maximum over all electrodes, regardless of positive lobes, was given by 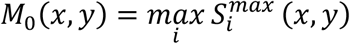.

## Acknowledgments

This work was supported by the Biotechnology and Biological Sciences Research Council (BBSRC, BH163322), Newcastle University Faculty of Medical Sciences and by the European Research Council (ERC) under the European Union’s Horizon 2020 research and innovation programme (grant agreement number 724822).

JdM, CT and ES designed the experiments; JdM, CT, DB, DBS, VK and ES performed the experiments; JdM, CT and ES analyzed the experimental data; FR and TG designed and performed the electrical imaging analysis; ES, CT and FR wrote the manuscript with input from the other authors.

